# Opposing Influence of Sensory and Motor Cortical Input on Striatal Circuitry and Choice Behavior

**DOI:** 10.1101/446708

**Authors:** Christian R. Lee, Alex J. Yonk, Joost Wiskerke, Kenneth G. Paradiso, James M. Tepper, David J. Margolis

## Abstract

The striatum is the main input nucleus of the basal ganglia and is a key site of sensorimotor integration. While the striatum receives extensive excitatory afferents from the cerebral cortex, the influence of different cortical areas on striatal circuitry and behavior is unknown. Here we find that corticostriatal inputs from whisker-related primary somatosensory (S1) and motor (M1) cortex differentially innervate projection neurons and interneurons in the dorsal striatum, and exert opposing effects on sensory-guided behavior. Optogenetic stimulation of S1-corticostriatal afferents in *ex vivo* recordings produced larger postsynaptic potentials in striatal parvalbumin (PV)-expressing interneurons than D1- or D2-expressing spiny projection neurons (SPNs), an effect not observed for M1-corticostriatal afferents. Critically, *in vivo* optogenetic stimulation of S1-corticostriatal afferents produced task-specific behavioral inhibition, which was bidirectionally modulated by striatal PV interneurons. Optogenetic stimulation of M1 afferents produced the opposite behavioral effect. Thus, our results suggest opposing roles for sensory and motor cortex in behavioral choice via distinct influences on striatal circuitry.

## Introduction

The basal ganglia comprise a circuit of interconnected nuclei that are involved in a variety of behavioral functions, including sensorimotor integration and the control of voluntary movement. The striatum is the largest input nucleus of the basal ganglia, and receives afferents from numerous areas of neocortex and thalamus [1,2]; [3–11]. In addition to promoting movement, activation of striatal neurons is also important in the termination of ongoing movement [12,13], as well as selecting an appropriate motor program and inhibiting competing motor programs [14]. The roles of sensory and motor cortical areas in the control of movement is a fundamental unresolved issue [15–17]. Primary sensory areas could play important roles through their corticostriatal projections, but this possibility has only recently begun to be explored [18–22]. Notably, the anterior dorsal striatum, which has been shown to be important for sensory-guided learning and behavior, receives overlapping projections from both sensory and motor cortical areas [3,4,8,9,23,24]

Excitatory afferents to the striatum innervate both classes of projection neurons known as D1- and D2-receptor expressing spiny projection neurons (SPNs), as well as an increasingly appreciated diversity of interneurons, including parvalbumin (PV)-expressing, GABAergic fast spiking interneurons [25–32]. Activation of D1-SPNs, constituting the direct pathway, tends to promote movement, while activation of D2-SPNs, constituting the indirect pathway, suppresses movement with both pathways exerting actions on other basal ganglia nuclei, and consequently on targets including the thalamus [33–36]. Conversely, striatal interneurons such as parvalbumin-expressing fast-spiking interneurons exert their effects locally by inhibiting both D1- and D2-SPNs as well as other striatal interneurons, which can have complex effects on behavior [23,24,36–41]. Although most of these neuron types receive excitatory input from numerous areas of neocortex [3,31], it remains unknown whether regionally distinct cortical areas differentially innervate striatal circuitry. It is possible that the balance between cortical targeting of SPNs versus interneurons is a key determinant of the behavioral effects of corticostriatal input, but this possibility has not been investigated.

Primary sensory areas of the neocortex have been extensively studied in the context of sensation and perception, but recent years have seen a greater appreciation for the role of sensory cortex in sensorimotor integration and motor control [17,42–44]. This is partly due to better understanding of the extensive interactions between sensory cortex, higher-order motor and association areas, and subcortical structures [45–47], as well as the ability to measure and manipulate neural activity in behaving subjects [48,49]. The primary somatosensory barrel cortex (S1) of the mouse vibrissal system is an important model system for investigating sensorimotor integration, including sensory-guided choice behavior [50–52]. After ascending sensory input arrives from sensory thalamus at L4 barrels, layer 2/3 and L5 excitatory projection neurons route information to multiple downstream cortical and subcortical targets. S1 projections to secondary somatosensory cortex (S2) and primary motor cortex (M1) have been well studied [52–58], but there is still little known about the circuitry or behavioral role of S1 projections to striatum. This is a major gap in knowledge because the S1-striatum projection is the largest anatomical projection from S1 [46]. We were specifically interested to investigate the anterior region of the dorsal striatum because, while S1 projects to both anterior and posterior striatum [17], the anterior region contains axonal projections from S1 and M1 [3,4], and thus may be important for sensorimotor integration.

We used *ex vivo* whole cell recordings of identified striatal SPNs and PV interneurons to measure the connectivity of both S1- and M1-corticostriatal inputs. We further interrogated the *in vivo* influence of S1- and M1-corticostriatal inputs to the same region of anterior dorsal striatum on tactile decision making. We report substantial differences in the effects of S1- and M1-corticostriatal inputs on striatal circuitry and sensory-guided behavior.

## Results

### Optogenetic activation of S1 or M1 corticostriatal afferents differentially engage striatal neurons

We used optogenetic activation of corticostriatal afferents from S1 or M1 with channelrhodopsin-2 (ChR2) combined with *ex vivo* whole cell current clamp recordings of identified striatal neurons to determine the striatal circuitry engaged by corticostriatal input from different sources (**Figure 1A**). We chose to investigate the anterior dorsal striatum because of prominent overlap between S1 and M1 inputs in this region (**Supplemental Figure 1**) [3,4]. Striatal D1-SPNs, D2-SPNs and PV interneurons were identified using tdTomato-expressing reporter mice and verified through their intrinsic properties in response to hyperpolarizing and depolarizing current injection (see Methods) (**Figure 1B-D**). We used current clamp recordings in the absence of inhibitory synaptic blockers, as opposed to voltage clamp recordings, to more closely mirror the natural physiological activity of striatal circuitry in response to S1 or M1 inputs. Activation of ChR2-expressing S1 corticostriatal afferents with a single, full-field light pulse (2.5 ms, 460-500 nm; through the 40x microscope objective) induced a depolarizing postsynaptic potential (PSP) in D1-SPNs, D2-SPNs and PV interneurons (**Figure 1E-G**). PSP amplitude was larger in PV interneurons (12.81 ± 2.37 mV, n = 9 neurons from 8 mice) compared to D1-SPNs (2.73 ± 0.71 mV, n = 17 neurons from 10 mice, p = 0.000008) or D2-SPNs (3.23 ± 0.95 mV, n = 10 neurons from 6 mice, p = 0.0001) (F_(2, 33)_ = 17.64, p = 0.000006; **Figure 1H**). Suprathreshold responses were sometimes encountered in PV interneurons, but only rarely in D1- or D2-SPNs. Measurable PSPs were found in all recorded neurons, suggesting that S1 innervates D1-, D2-SPNs and PV interneurons with high probability, but that PSP amplitude is biased toward PV interneurons.

**Figure 1:**
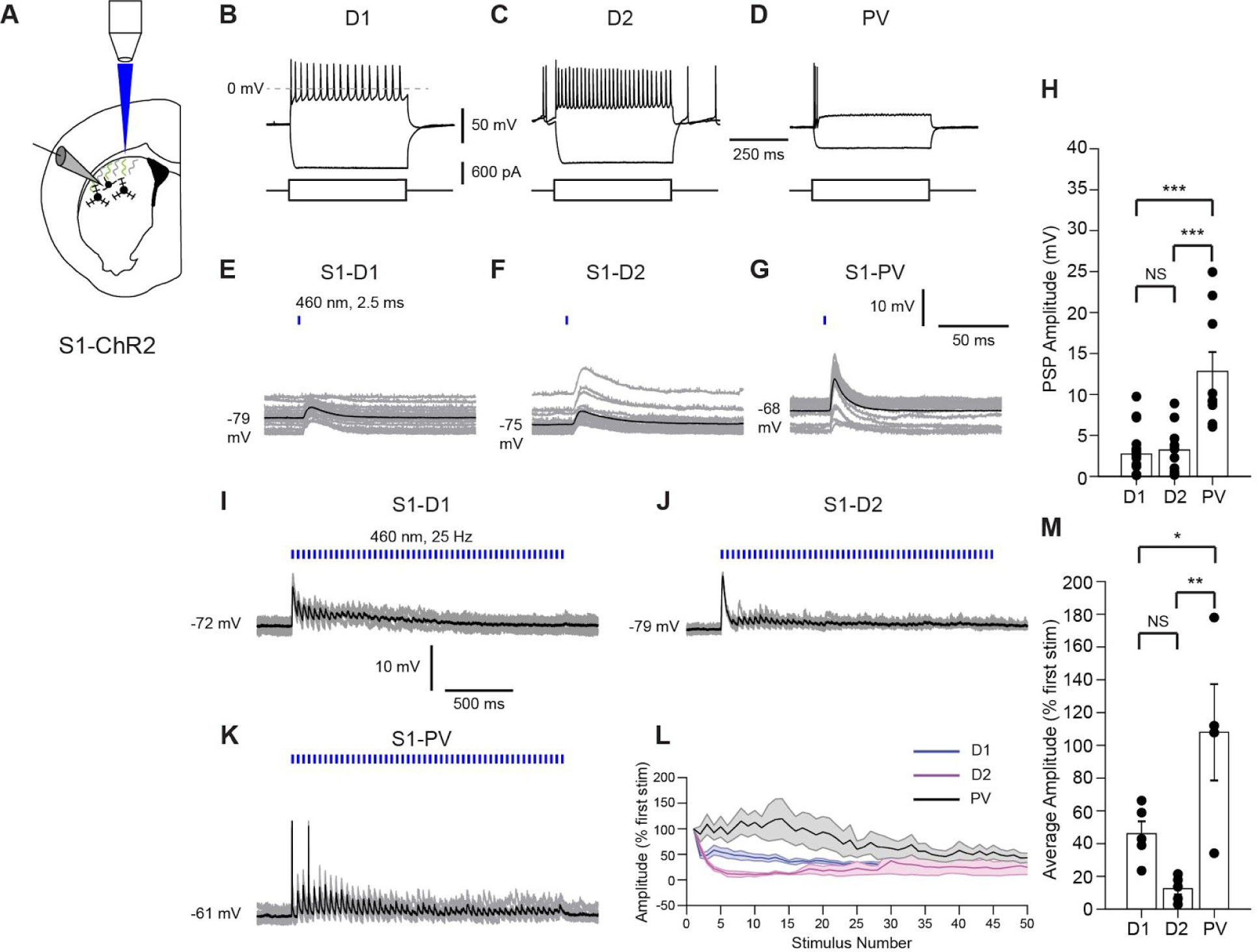
S1 corticostriatal input preferentially excites striatal PV interneurons. **(A)** Schematic diagram illustrating recording of striatal neurons and optogenetic activation of S1 corticostriatal afferents in an ex vivo slice of dorsal anterior striatum (1.4 to 0.4 mm anterior to bregma). See also Figure S1. Full field light was delivered through a 40x objective. **(B)** Characteristic physiological properties of D1- and **(C)** D2-SPNs and **(D)** PV interneurons in response to hyperpolarizing and depolarizing current injection. **(E)** Postsynaptic potentials (PSPs) evoked by optogenetic stimulation of S1 corticostriatal afferents in a D1-SPN. The average PSP is shown in black and individual traces in gray. Optogenetic stimulation is schematized by the blue line. **(F)** The postsynaptic potential evoked in a D2-SPN and **(G)** a PV interneuron. **(H)** The amplitude of the PSP evoked by optogenetic stimulation of S1 corticostriatal inputs was significantly larger in PV interneurons than in D1- or D2-SPNs. **(I)** Representative responses of a D1-SPN to optogenetic train stimulation of S1 corticostriatal inputs. Train stimulation is schematized by the blue lines. **(J)** The same is shown for a D2-SPN. Both D1- and D2-SPNs show strong short-term depression to train stimuli. **(K)** Representative response of a PV interneuron to S1 corticostriatal train stimulation. In contrast to the SPNs, train stimulation shows less synaptic depression through the train, and even some facilitation early on. Individual traces are shown in grey and the average in black. Truncated suprathreshold responses are shown, but were filtered to estimate the underlying PSP where necessary. **(L)** Summary plots of the PSP amplitude for each neuron type compared to first pulse amplitude. Mean is shown with SEM shaded. Both D1- and D2-SPNs show stronger synaptic depression during the train while the PV interneurons show a slight initial facilitation followed by depression later in the train. **(M)** Relative PSP amplitude (mean of pulses 5 to 14 compared to first pulse) is significantly greater in PV interneurons than in D1- or D2-SPNs. Data shown as mean ± SEM. ***p < 0.001, **p < 0.01, *p < 0.05, NS = not significant.

We further tested the synaptic dynamics of corticostriatal inputs during train stimulation. Train stimulation (2.5 ms pulses at 25 Hz for 2 s) of S1-corticostriatal afferents to D1-SPNs (n = 5 neurons from 4 mice) or D2-SPNs (n = 5 neurons from 3 mice) produced PSPs with strong synaptic depression, limiting the efficacy of the input during the train (**Figure 1I-J**). Conversely, when measured from PV interneurons (n = 4 neurons from 3 mice), train stimulation of S1 corticostriatal afferents produced PSPs with less depression (**Figure 1K**). Often, these PSPs exhibited slight facilitation early in the pulse train, consistent with previous findings [30]. When plotted as a percentage of the initial response amplitude, PSPs evoked in PV interneurons remained larger throughout the stimulus train (**Figure 1L**). The largest differences in PSP amplitudes between PV interneurons and D1- or D2-SPNs occurred during pulses 5 to 14, and were significantly different in this stimulus range (F_(2, 11)_ = 9.6, p=0.0039, PV vs D1 p = 0.0499, PV vs D2 p = 0.0034, D1 vs D2 p = 0.3963; **Figure 1L, M**).

We performed further *ex vivo* current clamp recordings to determine the effects of M1 input. In contrast to input from S1, optogenetic activation of M1-corticostriatal afferents evoked equally large amplitude PSPs in D1-SPNs (11.37 ± 2.25 mV, n = 13 neurons from 8 mice), D2-SPNs (13.35 ± 2.52 mV, n = 16 neurons from 10 mice) and PV interneurons (13.59 ± 3.07 mV, n = 13 neurons from 7 mice) (F_(2, 39)_ = 0.2, p = 0.82; **Figure 2A-H**). Suprathreshold responses were routinely encountered in all neuron types in response to stimulation of M1-corticostriatal input. Notably, PSP amplitudes recorded in PV interneurons in response to S1- or M1-corticostriatal afferent activation were indistinguishable (S1 12.81 ± 2.37, M1 13.59 ± 3.07 mV, p = 0.8554), suggesting that the observed differences in S1- and M1-evoked PSP amplitudes in D1- and D2-SPNs were not due to differences in cortical ChR2 expression.

**Figure 2:**
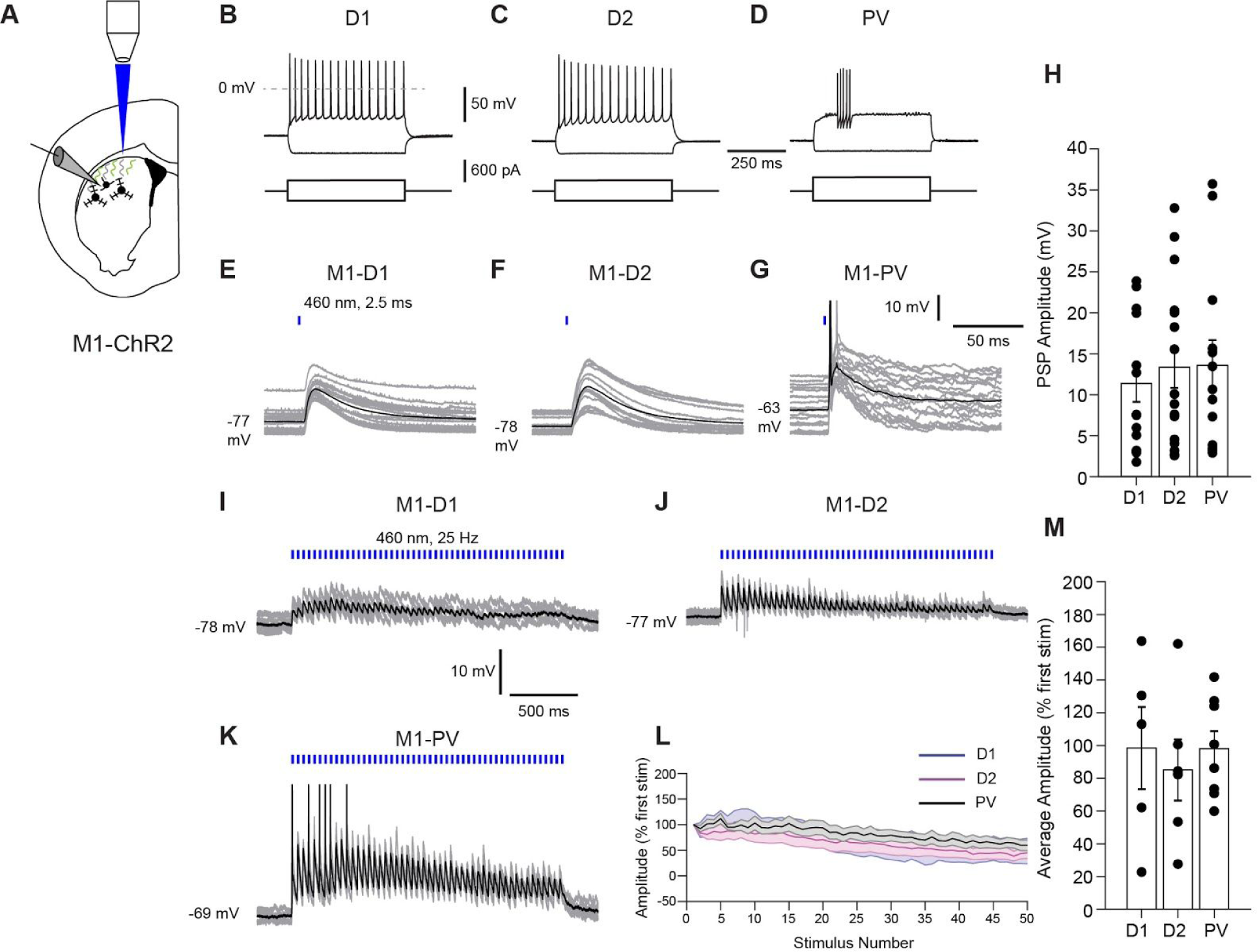
M1 corticostriatal input excites SPNs and PV interneurons equally. **(A)** Schematic diagram illustrating recording of striatal neurons and optogenetic activation of M1 corticostriatal afferents in an ex vivo slice. See also Figure S1. **(B)** Representative responses to hyperpolarizing and depolarizing current injection in D1-SPNs or **(C)** D2-SPNs as well as a **(D)** PV interneuron. **(E)** Postsynaptic potentials evoked by optogenetic stimulation of M1 corticostriatal afferents in a D1-SPN, **(F)** D2-SPN and **(G)** a PV interneuron. Average response shown in black and individual responses in grey. In some cases, action potentials have been truncated. Optogenetic stimulation is schematized by the blue line. **(H)** Amplitudes of PSPs evoked in each of these neuron types were similar. **(I)** Responses of a D1-SPN to train optogenetic stimulation of M1 corticostriatal afferents. **(J)** The same is shown for a D2-SPN and **(K)** a PV interneuron. Train stimulation is schematized by the blue lines. Truncated suprathreshold responses are shown, but were filtered and the underlying PSP estimated where necessary. **(L)** Stimulation of these afferents induced PSPs with similar levels of synaptic depression through the train. Data plotted as mean with shaded SEM. **(M)** When compared over the same range of stimulus pulses (5 to 14) that showed large deviations in PSP amplitude during stimulation of S1 corticostriatal input, PSPs evoked by M1 corticostriatal input did not differ in amplitude. Data presented as mean ± SEM.

Train stimulation of M1-corticostriatal afferents produced PSPs with minimal synaptic depression throughout the train in D1-SPNs (n = 5 neurons from 2 mice), D2-SPNs (n = 6 neurons from 4 mice), and PV interneurons (n = 8 neurons from 4 mice). PSP amplitude averaged over the 5 to 14 pulse range was not different among the three cell types (F_(2, 16)_ = 0.19, p = 0.8273) (**Figure 2I-M**), indicating that train stimulation of M1-corticostriatal afferents results in similarly small amounts of synaptic depression in both SPNs and PV interneurons. Overall, synaptic depression in SPNs was smaller for M1-compared to S1-corticostriatal afferents (M1 91.2 ± 14.6% n = 11 neurons, S1 29.3 ± 6.8% n = 10 neurons, t = 3.7075, p = 0.0015), but not PV interneurons (M1 98.0 ± 10.1% n = 8 neurons, S1 108.0 ± 29.4% n = 4 neurons, t = −0.3956, p = 0.7007).

Together, these results indicate that S1 provides larger amplitude synaptic input to striatal PV interneurons compared to either D1- or D2-SPNs. In contrast, the amplitudes of M1 synaptic inputs to D1-, D2-SPNs and PV interneurons were similar. These differences were further accentuated during train stimuli, as S1-corticostriatal input produced stronger synaptic depression in D1- and D2-SPNs compared to PV interneurons, an effect that did not occur for M1-corticostriatal input. Thus, a striking difference in S1-versus M1-corticostriatal innervation is the more efficacious innervation by S1 of PV interneurons compared to D1- and D2-SPNs.

### Input-specific effects on behavioral performance

To determine the effect of S1- or M1-corticostriatal input on sensorimotor behavior, we assessed how optogenetic stimulation of these regionally specific corticostriatal inputs affected performance of a whisker-dependent texture discrimination task in head-fixed mice (**Figure 3A-C**) [54,56,59]. In this Go/NoGo task, mice receive water reward for correct licking in response to the rough texture (Hit trial) and no water reward for licking in response to the smooth texture (False Alarm trial). Textures are presented to the whiskers via a motorized stage. Light was delivered to the same region of the anterior dorsal striatum as used for electrophysiology experiments (above) through an implanted optical fiber (**Supplemental Figure 1**). We measured the effects of optogenetic stimulation of S1- or M1-corticostriatal afferents on changes in Hit Rate, False Alarm Rate, Sensitivity, and Bias compared to the start of the session. Mice completed 50 trials without optogenetic stimulation followed by 77 trials with stimulation. Optogenetic stimulation of S1-corticostriatal input led to reduction in behavioral responding, as evident in a raster plot of licking in a representative session before and during stimulation (**Figure 3D**). As a result of S1-corticostriatal stimulation, we found significant reductions in Hit Rate (0.62 ± 0.08 to 0.39 ± 0.11, p = 0.005, n = 6), FA Rate (0.39 ± 0.10 to 0.21 ± 0.06, p = 0.013, n = 6), and Bias (−0.04 ± 0.31 to −0.78 ± 0.35, p = 0.011, n = 6), while Sensitivity (d’) decreased, but not significantly (d’: 0.93 ± 0.34 to 0.62 ± 0.40, p = 0.245, n = 6) (**Figure 3E**).

**Figure 3:**
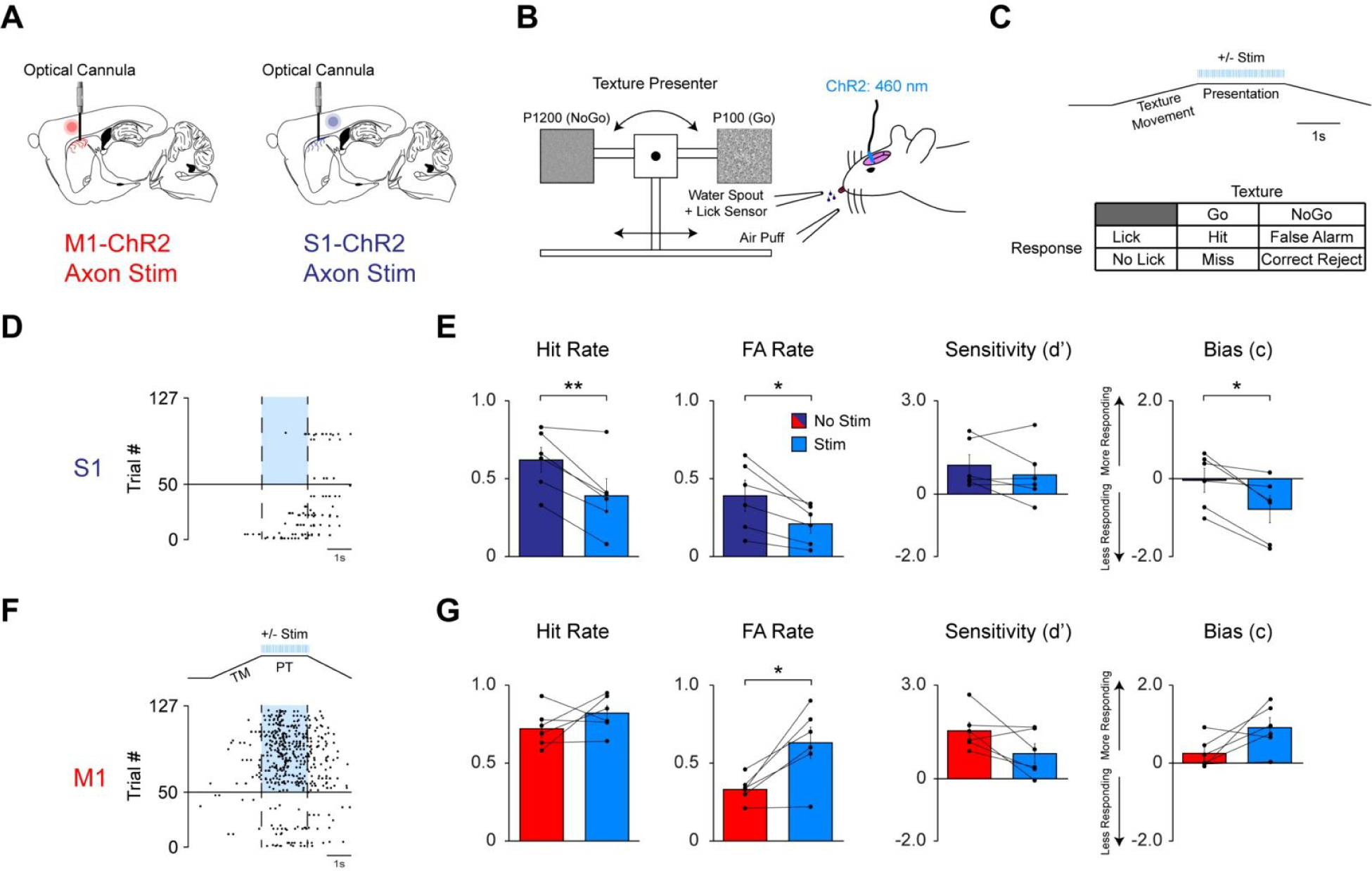
S1 corticostriatal input inhibits responding while M1 corticostriatal input promotes responding in a tactile discrimination task. **(A)** Schematic sagittal view (+1.8 mm lateral from midline) showing optical cannula placement along with AAV (ChR2) transfected M1 (*Left)* and S1 (*Right)* terminals in the striatum (adapted from [74]). The red and purple circle illustrates the M1 and S1 injection sites, respectively. **(B)** Schematic of optogenetic stimulation during tactile discrimination task. **(C)** *Top:* Representation of trial structure and stimulation timing. *Bottom:* Possible outcomes for each stimulus-response pair. **(D)** Raster plot showing lick responses (black dots) throughout a single representative session during baseline and S1 corticostriatal stimulation. Vertical dashed lines indicate the start and end of the presentation time, during which mice can respond. The horizontal black line indicates the onset of optical stimulation at trial 51. The blue portion between the dashed lines indicates the timing of optical stimulation. **(E)** Effects of S1 corticostriatal stimulation (n = 6 mice) on Hit Rate, FA Rate, Sensitivity and Bias. Data are mean (± SEM) for baseline and stimulated conditions. **(F)** Raster plot showing lick responses (black dots) throughout a representative session during baseline and M1 corticostriatal stimulation. Vertical dashed lines indicate the start and end of the presentation time. The horizontal black line indicates the onset of optical stimulation at trial 51. The blue portion between dashed lines indicates the timing of optical stimulation. Trial structure and ChR2 stimulation during the response window is schematized above. **(G)** Effects of M1 corticostriatal stimulation (n = 6 mice) on Hit Rate, FA Rate, Sensitivity, and Bias. Data are mean (± SEM) for baseline and stimulated conditions. Hit Rate is the proportion of successful Go trials divided by the total number of Go trials. FA Rate is the proportion of unsuccessful NoGo trials divided by the total number of NoGo trials. Sensitivity is the capacity to discriminate between the Go and NoGo texture. Bias is the overall responding during the session. Effects of optogenetic manipulations are specific to the task and the task is dependent on tactile input (see also Figure S2 and S3). \**p* < 0.05, \*\**p* < 0.01.

The behavioral effects of optogenetic stimulation of M1-corticostriatal input were markedly different, leading instead to increased behavioral responding, as seen in the raster plot of licking in a representative behavioral session (**Figure 3F**). M1-corticostriatal stimulation led to a significant increase in FA Rate (0.33 ± 0.04 to 0.63 ± 0.10, p = 0.013, n = 6; **Figure 3G**) reflecting an overall increase in responding. Hit Rate (0.72 ± 0.06 to 0.82 ± 0.05, p = 0.285, n = 6), Sensitivity (d’: 1.53 ± 0.28 to 0.80 ± 0.32, p = 0.086, n = 6), and Bias (0.24 ± 0.17 to 0.90 ± 0.26, p = 0.071, n = 6) did not change significantly (**Figure 3G**). Comparison of S1- and M1-corticostriatal data showed that the overall effects of stimulation were highly divergent for Bias (M1: 0.90 ± 0.26, S1: −0.78 ± 0.35, p = 0.0018, n = 6 for each condition), but not Sensitivity (M1: 0.80 ± 0.32, S1: 0.62 ± 0.40, p = 0.709, n = 6 for each condition).

To determine whether optogenetic activation of S1- or M1-corticostriatal input affects licking behavior directly, we delivered optogenetic stimulation during a cued-reward version of the task to the same subjects. In this case, textures were not presented and water reward was delivered automatically in each trial following a cue tone and audible solenoid click. Neither S1-nor M1-corticostriatal stimulation affected lick rate during this task (S1: 4.23 ± 1.46 to 4.87 ± 1.65 licks/second, p = 0.1063; M1: 3.30 ± 1.02 to 2.67 ± 0.85 licks/second, p = 0.1016, n = 3 mice for each group), suggesting that the effects of S1- or M1-corticostriatal activation that we observed during the texture discrimination task are not due to effects on licking behavior.

We performed further behavioral tests in the same subjects to determine whether the effects of S1- and M1-corticostriatal stimulation were specific to the texture discrimination task by stimulating S1 and M1 corticostriatal afferents during open field (OF) exploration and while testing mice on an accelerating rotarod (RR). We observed no overall effect on locomotion during open field exploration or during rotarod testing, even with prolonged stimulation for 2 minutes during open field exploration or up to 5 minutes during rotarod performance compared to baseline performance (OF: M1: p = 0.498, n = 6; S1: p = 0.505, n = 5; PV: p = 0.101, n = 5; RR: M1: p = 0.105, n = 5; S1: p = 0.087, n = 5; PV: p = 0.573, n = 6; **Supplemental Figure 2**), suggesting that activation of specific corticostriatal inputs affects behavior in a more subtle way than strong, direct stimulation of SPNs which can induce changes in locomotion and other behavior [35,41]. Finally, testing mice on the texture discrimination task after trimming the whiskers showed significant reduction of both Bias and Sensitivity, demonstrating that the task is dependent on whisker-based tactile input (**Supplemental Figure 3**).

Overall, the results of decreased behavioral responding during S1-corticostriatal stimulation and increased behavioral responding during M1-corticostriatal stimulation indicate opposing effects of S1- and M1-corticostriatal activation on sensory-guided behavior.

### Optogenetic stimulation of striatal PV interneurons suppresses responding in the tactile discrimination task

How do S1- and M1-corticostriatal inputs lead to opposite effects on behavior? We hypothesized that the biased innervation of PV interneurons by S1 (above) could play a key role. We tested whether direct optogenetic activation of PV interneurons caused a similar behavioral effect as S1 optical stimulation (**Figure 4A-C**). An optical fiber was implanted above dorsal striatum in mice expressing ChR2 in PV interneurons, and mice were trained on the task as above. Mice completed 50 trials without optogenetic activation followed by 77 trials with activation of PV interneurons. Optogenetic activation of striatal PV interneurons led to a reduction of behavioral responding, as shown in a representative session (**Figure 4D**). Overall, Hit Rate (0.60 ± 0.08 to 0.42 ± 0.12, p = 0.007, n = 6), FA Rate (0.29 ± 0.03 to 0.19 ± 0.04, p = 0.01, n = 6), and Bias (−0.18 ± 0.18 to −0.72 ± 0.34, p = 0.03, n = 6; **Figure 4E**) decreased significantly, while Sensitivity (d’: 1.00 ± 0.37 to 0.57 ± 0.50, p = 0.053, n = 6; **Figure 4E**) decreased, but not significantly. Thus, direct activation of PV interneurons exerts a suppressive influence on behavioral responding, similar to S1-corticostriatal stimulation.

**Figure 4:**
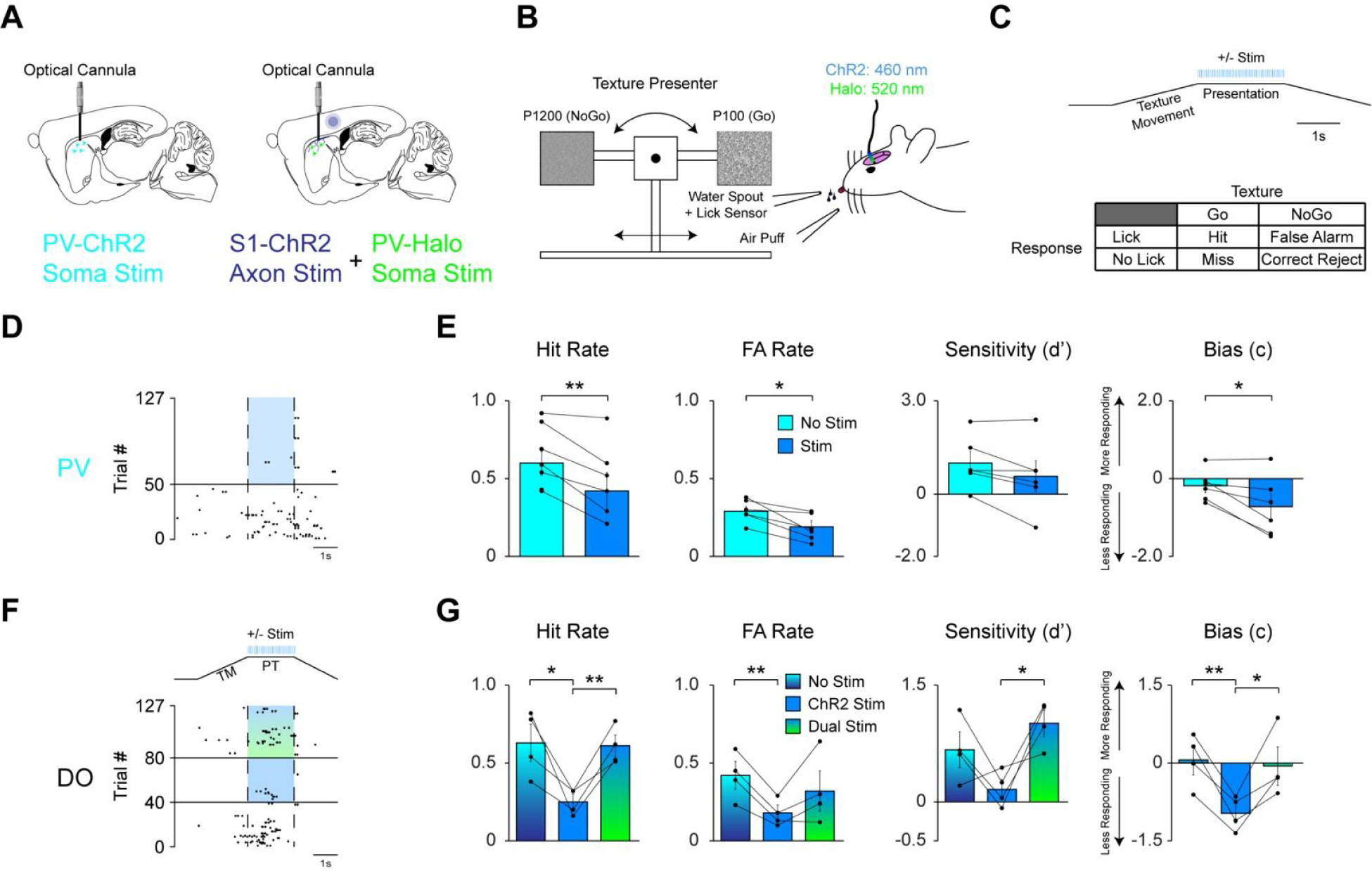
Response inhibition caused by S1 corticostriatal input is mediated by striatal PV interneurons. **(A)** Schematic sagittal view (+1.8mm lateral from midline) show optical cannula placement along with transgenic PV interneuron somas (*Left*) and AAV (S1: ChR2; PV: NpHR) transfected S1 terminals and PV interneuron somas (*Right*) in the striatum (adapted from [74]). The purple circle illustrates the S1 injection site. **(B)** Schematic of optogenetic stimulation during tactile discrimination task. In some experiments, PV interneurons expressing ChR2 were directly stimulated with blue light (460 nm). In other experiments, PV interneurons were inhibited by activating NpHR with green light (520 nm), while S1 terminals were simultaneously stimulated with blue light (460 nm). **(C)** *Top:* Representation of trial structure and train stimulation timing. *Bottom:* Possible outcomes for each stimulus-response pair. **(D)** Raster plot showing lick responses (black dots) throughout a representative session during baseline and direct activation of PV interneurons expressing ChR2. Vertical dashed lines indicate the start and end of the presentation window. The horizontal black line indicates the onset of optical stimulation at trial 51. The blue portion between the dashed lines indicates the timing of optical stimulation. **(E)** Effects of direct PV interneuron stimulation (n = 6 mice) on Hit Rate, FA Rate, Sensitivity, and Bias. Data are mean (± SEM) for baseline and stimulated conditions. **(F)** Raster plot of lick responses throughout a representative session during baseline, S1 corticostriatal stimulation (ChR2) only, and simultaneous S1 corticostriatal stimulation (ChR2) and optogenetic suppression of PV interneurons (NpHR). Vertical dashed lines indicate the start and end of the presentation window. The horizontal black line at trial 41 indicates the onset of S1 corticostriatal stimulation. The horizontal black line at trial 81 indicates the onset of simultaneous optogenetic suppression of PV interneurons with NpHR, as indicated by the blue/green gradient fill. Trial structure and train stimulation of ChR2, as well as tonic activation of NpHR, during the response window is schematized above. **(G)** Effects of S1 corticostriatal stimulation and simultaneous S1 corticostriatal stimulation and PV interneuron suppression (n = 4 mice) on Hit Rate, FA Rate, Sensitivity, and Bias. Data are mean (± SEM) for baseline, S1 only, and dual optogenetic stimulation. Effects of optogenetic manipulations are specific to the task and the task is dependent on tactile input (see also Figure S2 and S3). * *p*< 0.05, ** *p* < 0.01.

Further, we delivered optogenetic activation during a cued-reward version of the task, when textures were not presented, but water reward was delivered automatically in each trial following a cue tone and audible solenoid click. Optogenetic activation of PV interneurons had no effect on lick rate during this cued reward task indicating no direct effect of PV neuron excitation on licking (1.86 ± 0.57 to 1.69 ± 0.84 licks/second, p = 0.6403; n = 3 mice). We observed a slight but significant decrease in locomotion during open field exploration during the first epoch of stimulation, with no change in locomotion during later 2 minute epochs of stimulation. Rotarod performance was unaffected by stimulation up to 5 minutes (**Supplemental Figure 2**). Overall, direct activation of PV interneurons in dorsal striatum had clear effects on texture discrimination, but minimal effects on other behaviors, including licking, open field and rotarod.

### PV cell suppression abolishes inhibitory effects of S1 stimulation on behavior

Our results thus far suggest that striatal PV interneurons mediate the suppressive effects of S1 on behavior. We next tested whether inhibiting these interneurons could attenuate or reverse the behavioral inhibition produced by S1 stimulation. We tested this in mice coexpressing halorhodopsin (NpHR) in striatal PV interneurons and ChR2 in S1 corticostriatal afferents. Mice were again trained on the texture discrimination task. Mice performed 40 baseline trials, followed by 40 S1-ChR2 optical stimulation trials (460 nm), followed by 47 trials with S1-ChR2 stimulation and concurrent inhibition of PV interneurons with NpHR (460 plus 520 nm). We found that optogenetic inhibition of PV interneurons could reverse the suppressive effect of S1-corticostriatal stimulation on behavior, as shown in a representative session (**Figure 4F**) and reflected in significant differences for all behavioral parameters among the three conditions (repeated measures ANOVA: Hit Rate (F_(2, 6)_ = 9.66, p = 0.0133, n = 4), FA Rate (F_(2, 6)_= 7.08, p = 0.0264, n = 4), Sensitivity (F_(2, 6)_ = 8.18, p = 0.0193, n = 4), and Bias (d’: F_(2, 6)_ = 10.6, p = 0.0107, n = 4) (**Figure 4G**). Paired contrasts revealed a significant decrease in Hit Rate following optogenetic stimulation of S1 corticostriatal afferents, as expected (p = 0.0426). Strikingly, Hit Rate increased significantly following simultaneous stimulation of S1 corticostriatal afferents and optogenetic inhibition of PV interneurons (p = 0.0088) returning to control levels (baseline vs. dual stim, p = 0.8353). Similarly, there was a significant decrease in FA Rate following S1 corticostriatal stimulation (p = 0.0078) which was lost with simultaneous inhibition of PV interneurons (baseline vs. dual stim, p = 0.3105). Changes in Sensitivity and Bias induced by S1 corticostriatal stimulation were also reversed by simultaneous inhibition of PV interneurons (**Figure 4G**). Sensitivity increased significantly following simultaneous S1 corticostriatal stimulation and inhibition of PV interneurons compared to stimulation of S1 input alone (p = 0.0375, baseline vs. dual opto, p = 0.0559). Bias decreased significantly following S1 corticostriatal stimulation (p = 0.0095) as expected, and returned to control levels during simultaneous inhibition of PV interneurons (S1 stim vs dual opto p = 0.0361, baseline vs. dual opto, p = 0.6936).

Thus, our results indicate that S1-corticostriatal input exerts an inhibitory influence on sensory-guided behavior, opposite the effects of M1-corticostriatal input. The behavioral inhibition could be reproduced by direct striatal PV interneuron activation, and furthermore could be restored by suppression of PV interneurons during S1-corticostriatal activation. Our results suggest a novel influence of S1 on corticostriatal circuitry and behavior, and implicate striatal PV interneurons as key players in this process.

## Discussion

We have shown that corticostriatal inputs from sensory and motor cortical areas differentially engage neurons in the dorsal striatum, leading to distinct influences on behavior. Specifically, S1-corticostriatal input primarily activates PV interneurons and suppresses responding in a tactile discrimination task, while M1-corticostriatal input activates SPNs and PV interneurons equally, and produces increased responding in the task. Bidirectional optogenetic manipulations also revealed a central role for striatal PV interneurons in behavioral control within a tactile discrimination task but not in overall movement or motivated licking. These are the first data to show that differential activation of striatal neurons by discrete neocortical areas leads to opposite effects on sensory-guided behavior.

### The role of the basal ganglia in response inhibition

For nearly three decades, it has been appreciated that the basal ganglia are involved in both the promotion of movement through activation of direct pathway SPNs and the suppression of movement through activation of indirect pathway SPNs [33,34]. Natural movements are complex and require the activation of some motor programs and the suppression of others, resulting in the activation of both pathways during behavior [14,36,60]. However, the movement promoting influence of the direct pathway and the movement suppressing influence of the indirect pathway is supported by the bulk of available evidence [35,36]. The mechanisms that could lead to activation of one striatal pathway over the other have remained incompletely understood, though the demonstrated role of the hyperdirect pathway from cortex to the subthalamic nucleus of the indirect pathway in stopping ongoing movements has strengthened the idea that differential activation of the direct and indirect pathways can bias behavior [61,62]. Here we demonstrate an additional pathway for action suppression through S1-corticostriatal input that preferentially activates PV interneurons. An earlier study using electrical stimulation of primate sensory cortex and detection of immediate early gene expression in striatum also suggested a preferential activation of striatal PV interneurons by S1 activation [63]. It appears increasingly plausible that S1 is an important mediator of behavioral control through its actions on striatal PV interneurons.

### Sensory and motor corticostriatal signaling

S1 barrel cortex has long been recognized as a key locus for whisker-guided behaviors. While S1 is densely interconnected with many cortical and subcortical areas, including thalamus, higher-order sensory, motor, and association cortices [45–47], the detailed circuitry and behavioral roles of these pathways are just beginning to be understood [64,65]. For example, S1 interactions with M1 and S2 are now recognized as important for detection of sensory stimuli, behavioral performance, and learning [53–56,58,66,67]. Some of the largest anatomical projections from both S1 and M1 are to the striatum, where S1 and M1 projection fields overlap in the anterior portion of the dorsal striatum [3,4,46,68,69]. This overlap even occurs at the level of individual SPNs and PV interneurons which have been shown to receive convergent corticostriatal input from S1 and M1 [4,27]. Remarkably, in spite of the potential importance of corticostriatal signaling for sensorimotor function, the circuitry and behavioral impact of S1- and M1-corticostriatal projections have remained unresolved. *In vivo* recordings have found that both D1- and D2-SPNs receive whisker-driven synaptic input [19,20], but differences between S1 and M1 input, and potential recruitment of identified interneurons was not investigated.

We targeted our investigation to the dorsal anterior striatum, based on recent evidence that this region is involved in sensory-guided learning and behavior [23,24]. We found that activation of S1-corticostriatal input more strongly innervates striatal PV interneurons than either D1- or D2-SPNs and leads to behavioral inhibition, whereas activation of M1-corticostriatal input equally innervates PV interneurons and SPNs, and leads to behavioral activation. These results represent a previously unappreciated difference between S1- and M1-corticostriatal inputs and have implications for understanding how the striatum integrates inputs from diverse cortical and subcortical sources to produce behavioral responses. Activity in posterior striatum has recently been associated with auditory guided behavior [22]. The extent to which the prominent S1 input to posterior striatum [3,17] is also involved in tactile-guided behavior is still unknown, and would be an important topic for future studies.

### Influence of striatal PV interneurons on circuitry and behavior

A key finding of our study is that direct activation of striatal PV interneurons suppresses responding in the tactile discrimination task in a similar fashion to activation of S1-corticostriatal afferents. These results are largely consistent with previous studies showing that optogenetic activation of striatal PV interneurons can disrupt behavior in lever press and olfactory learning tasks [24,36]. Striatal PV interneurons are powerfully activated by cortical inputs and provide feedforward GABAergic inhibition to SPNs [70–72]. One mechanism that might underlie the suppressive effect of PV interneuron activation on responding is the slight preference for PV interneurons to innervate direct pathway SPNs [39]. It is plausible that corticostriatal input from S1 leads to preferential inhibition of D1-expressing striatonigral SPNs, thus disinhibiting the inhibitory output of the basal ganglia from the substantia nigra pars reticulata/internal globus pallidus/entopeduncular nucleus and suppressing movement. Our data showing stronger short-term synaptic depression for S1 inputs to SPNs compared to PV interneurons, whether monosynaptic or polysynaptic, could further serve to accentuate the bias toward recruitment of PV interneurons during repetitive cortical activation.

Furthermore, we found that optogenetic inhibition of PV interneurons removes the suppressive effect of S1 corticostriatal input on task performance. This is important for two reasons. First, this result provides evidence that S1-corticostriatal input operates via PV interneurons. Second, it also argues against possible antidromic activation of S1 cell bodies and other downstream projections as a result of optogenetic activation of corticostriatal terminals. If antidromic activation did occur, it would still have occurred during simultaneous inhibition of striatal PV interneurons, yet no behavioral inhibition resulted (**Figure 4**). Instead, mice performed the task at baseline levels under these conditions, providing further evidence against the possibility that S1-corticostriatal terminal stimulation caused interference of behavior-related sensory processing.

Overall, our data suggest that PV interneuron activation, whether directly or through activation of S1 corticostriatal input, suppresses task-relevant sensorimotor behavior while leaving other movements intact. Understanding the detailed anatomical and biophysical mechanisms underlying differences in synaptic strength, and the engagement of striatal microcircuitry [2,37,73] during behavioral performance, will be important areas for future work.

## Conclusions

We have shown that input from primary sensory cortex (S1) preferentially engages striatal PV interneurons in the striatum and suppresses responding in a tactile discrimination task. Conversely, corticostriatal input from primary motor cortex (M1) engages D1- and D2-SPNs as well as PV interneurons, and promotes responding in the task. Our findings suggest that corticostriatal input from regionally and functionally specific areas of cortex can have different physiological and behavioral consequences. It will be important in future studies to investigate the dynamics of S1- and M1-corticostriatal neuronal activity during behavior, and to determine other potential differences in striatal circuit innervation by diverse sources.

## Acknowledgements

The authors thank Battulga Amarbat and Neeha Pathan for assistance with behavioral experiments, Ben Pasternak and Wahhaj Khokhar for assistance with analyzing lick data, and Dr. Mark West for providing D1- and D2-cre mice used for some experiments. This work was supported by grants from the Rutgers Brain Health Institute Pilot Grant Program to DJM and JMT, and the National Institutes of Health (R01NS094450, DJM; R01NS034865, JMT).

## Author Contributions

CRL, AJY, JW, KGP, JMT, and DJM designed research. CRL, AJY, and JW performed research. CRL, AJY, and JW analyzed data. CRL, AJY, and DJM wrote the manuscript. All authors approved the final version of the manuscript.

## Experimental Model and Subject Details

All procedures were approved by the Rutgers University Institutional Animal Care and Use Committee (IACUC; protocol #: 13-033). All mice used in experiments were housed in a reverse light cycle room (lights off from 08:00 to 20:00) with food and water available *ad libitum*, unless subjected to water restriction during behavioral training. Both male and female mice were used for electrophysiological and behavioral experiments and were adult at the time of the first surgical procedures (average 7.1 weeks, range 4.1 to 13.4 weeks). For electrophysiological experiments D1- and D2-SPNs as well as PV-interneurons were identified by expressing tdTomato in those neuron types using B6.Cg-Tg(Drd1a-tdTomato)6Calak/J (The Jackson Laboratory) [75] or one of the following mice crossed with Ai14 (The Jackson Laboratory) [76]: B6.FVB(Cg)-Tg(DrD1cre)EY262Gsat/Mmucd(Gensat),B6.FVB(Cg)-Tg(DrD2cre) ER44Gsat/Mmucd (Gensat), or B6.129P2-Pvalbtm1(cre)Arbr/J [29,77]. Mice used for behavioral experiments were often litter mates of mice used for electrophysiology experiments, but lacked a transgenic allele. In some cases B6.129P2-Pvalbtm1(cre)Arbr/J mice crossed with Ai14 (The Jackson Laboratory), or if ChR2 was expressed in PV interneurons, Ai32 (The Jackson Laboratory) were used for behavior [69,76]. In mice were NpHr was virally expressed in PV interneurons, B6.129P2-Pvalbtm1(cre)Arbr/J (The Jackson Laboratory) mice were used [77]. During behavioral testing, daily water intake was restricted to ~1.5 mL per mouse per day. This was done to motivate performance of the behavioral task described below. Baseline body weight was measured once prior to water restriction and daily thereafter. Body weight decreased on average to 84.24 ± 0.81% of their original weight, consistent with levels of restriction used to motivate behavior [78]. All handling and behavioral experiments were conducted during the dark phase of the cycle.

## Method Details

### Viral Vectors and Stereotaxic Injection

Unilateral injections of AAV1-CaMKIIa-hChR2(H134R)-eYFP.WPRE.hGH (Penn Vector Core; Addgene 26969P [79]) targeted either left whisker primary sensory cortex (S1) or left whisker primary motor cortex (M1). Briefly, mice were anesthetized with isoflurane (4% induction, 0.8-1.5% maintenance) and placed onto a stereotaxic frame (Stoelting) that had a feedback controlled heating blanket maintained at 36°C (FHC) on the base. The scalp was cleaned with Betadine (Purdue Products) followed by 70% ethanol (Fisher) three times. Ketorolac (5 mg/kg) (Hospira) and Bupivacaine (0.25%) (0.1 mL, Fresenius Kabi) were injected subcutaneously into the left flank and scalp, respectively. A midline incision was made and the skull was exposed. The skull was leveled in the dorsoventral plane by ensuring equal bregma and lambda coordinates. For M1 and S1 injections, a craniotomy was made at the following coordinates (S1 coordinates with respect to bregma: AP = −1.0 mm; ML = +3.3 mm; DV = −0.6 mm; M1 coordinates with respect to bregma: anteroposterior (AP) = +1.6 mm; mediolateral (ML) = +1.5 mm; dorsoventral (DV) = −0.6 mm). In mice where NpHR was expressed in PV interneurons, AAV5-Ef1a-DIO eNpHR 3.0-EYFP was injected into striatum (addgene 26966 [80]) at the following coordinates (AP) = 1 mm; (ML) = 1.8 mm (DV) = −2.0 mm in B6.129P2-Pvalbtm1 (cre)Arbr/J mice (The Jackson Laboratory) [77]. The micropipette was slowly lowered to the proper depth and allowed to sit for 5 minutes. Following this, 210 nL of ChR2 virus solution or 280 nl of NpHR virus solution, diluted 1:1 with phosphate buffered saline was pressure injected over 5 minutes followed by a delay of an additional 5 minutes to allow for viral diffusion into the tissue. The micropipette was then slowly raised. The scalp was closed and secured with silk sutures and tissue glue. After surgery, mice were placed in clean, temporary housing and monitored for 72 hours. After this monitoring period, mice were transferred to their home cages and allowed to recover for at least 4 weeks before either electrophysiological or behavioral experiments, permitting viral expression in the M1 and S1 axon terminals located in the striatum.

### *Ex vivo* whole-cell current clamp recordings

Mice were induced with isoflurane (3%) and deeply anesthetized with ketamine and xylazine (300/30 mg/kg, i.p.). Mice were then transcardially perfused with either cold or room temperature modified ACSF containing (in mM) NMDG 103, KCl 2.5, NaH_2_PO_4_ 1.2, NaHCO_3_ 30, HEPES 20, glucose 25, HCl 101, MgSO_4_ 10 which was continuously bubbled with 95% O_2_ 5% CO_2_. Coronal slices (300 μm) were cut on a Leica VT1200S vibratome. Following a short recovery in the same solution at 36°C, slices were transferred to ACSF which contained (in mM) NaCl 124, KCl 2.5, NaHCO_3_ 26, NaH_2_PO_4_ 1.2, glucose 10, pyruvate 3, MgCl_2_ 1, and CaCl_2_ 2 which was continuously bubbled with 95% O_2_ and 5% CO_2_ for at least one hour before recording. Recordings were obtained in the same solution. The pipette solution contained K methanesulfonate 130, KCl 10, HEPES 10, MgCl_2_ 2, Na_2_ATP 4, and Na_2_GTP 0.4 at pH 7.25 and 290-295 mOsm/L. Patch pipettes (2-5 MΩ) were constructed from 2.0 mm o.d. borosilicate glass (Warner Instruments) pulled using a Sutter P-1000 horizontal puller. Current clamp recordings were obtained from identified neuron types using mice expressing tdTomato in D1-, D2-, or PV-expressing neurons. When recording from SPNs, unlabeled neurons were assumed to belong to the other projection neuron type when the neuron exhibited characteristic physiological properties of SPNs such as little to no sag in voltage response after hyperpolarizing current injection and a ramp depolarization and regular action potential firing in response to depolarizing current injection [26]. Rarely, unlabeled neurons could be identified as a fast spiking interneuron based on physiological characteristics, the most prominent of which is a bursty action potential pattern in response to depolarizing current injection [26]. Neurons were recorded in the dorsal anterior striatum (approximately 1.4 to 0.4 mm anterior to bregma) which receives input from both S1 and M1 [3,4,27]. Channelrhodopsin-2 (ChR2) was activated by illumination with a 2.5 ms, 460-500 nm LED light pulse (1.2 mW measured after the objective, Thorlabs) delivered through the objective lens (40x) of an Olympus BX51WI microscope. Stimulation was presented once every 30 seconds for 20 sweeps. Slices were submerged and continuously superfused with ACSF during recording. Data were acquired with a HEKA EPC10 amplifier and digitized at 20 kHz in Patchmaster. Voltages reported in the example traces have not been corrected for the liquid junction potential.

Analysis of physiology data was carried out using custom scripts written in MATLAB. Single pulse postsynaptic potential amplitudes (PSPs) were calculated from averaged sweeps of up to 20 stimulations and were defined as the difference between the peak amplitude achieved within 500 ms centered around the time of optogenetic stimulation and the baseline membrane potential which itself was calculated as the average membrane potential between 1 s and 500 ms prior to optogenetic stimulation. Suprathreshold responses were excluded from analysis by removing sweeps containing action potentials from the average if only a small number of sweeps had action potentials in them or by estimating the size of the subthreshold PSP by filtering the data with a lowpass butterworth filter with a cutoff frequency of between 30 and 50 Hz. Traces were also excluded if the resting membrane potential drifted substantially as identified by visual inspection. Note that we use the term PSP instead of EPSP (excitatory PSP) because EPSPs were not isolated under our recording conditions. We chose this design to mimic in vivo conditions, where cortical input can activate monosynaptic excitation as well as disynaptic feedforward inhibition.

Train stimuli consisted of 2.5 ms pulses of light delivered at 25 Hz every 30 seconds for 5 sweeps. Sweeps were averaged and the amplitude of the PSPs calculated. Suprathreshold events were filtered with a lowpass butterworth filter with a cutoff frequency of 50 Hz. For each pulse interval, if the amplitude of the PSP was greater than 1.5 times the amplitude of the previous PSP, as would be the case in a suprathreshold response, filtered data was used to estimate the underlying PSP. If the first stimulation contained a suprathreshold response, filtered data was used for that pulse. The amplitude of the first PSP was calculated as 100% and amplitudes of subsequent PSPs in the train were calculated as a percentage of the first PSP.

### Optical Cannula Implantation Surgery

Mice designated for behavior testing were allowed to recover for three weeks prior to the fiber optic implant surgery. Following the implant surgery, mice were allowed to recover for an additional week prior to handling. A custom head post was fitted to each mouse using methods similar to those previously described [54,56,59,81]. Briefly, mice were anesthetized with isoflurane (4% induction, 0.8-1.5% maintenance) and mounted onto a stereotaxic frame (Stoelting) with a feedback controlled heating blanket maintained at 36°C (FHC) on the base. After the scalp was reflected, connective tissue was removed from the skull by gentle scraping and a blue light curable bonding agent (iBond, Heraeus Kulzer) was applied to the skull followed by a ring of dental composite around the outer edge (Charisma, Kulzer). A craniotomy was made above the left striatum for optical cannula insertion (coordinates with respect to bregma: AP = +1.0 mm; ML = +1.8 mm; DV = −2.0 mm). The optical cannula (length cut to 3 mm, 0.50 nA; 200 µm diameter; Thorlabs) was secured with dental composite (Tetric Evoflow, Ivoclar Vivadent). A custom aluminum head post (weight, < 1 g) was also cemented to the contralateral hemisphere. The scalp was closed around the resulting head cap using silk sutures and tissue glue.

After surgery, mice were allowed to recover in their home cages for one week before any handling was performed. Following the post-surgical recovery period, mice were handled daily for at least one week. During this handling period, mice were also acclimated to head fixation. This was done by placing them within a tube (length 14 cm; inner diameter 3.5 cm) attached to a custom platform (length 16.75 cm; width 12.25 cm). The platform contained an aluminum crossbar with screw holes to mount the head post and secure the mouse’s head.

### *In Vivo* Optogenetics

Two high-powered LEDs (ChR2: 460 nm; NpHR: 520 nm; Prizmatix) and LED current drivers (Prizmatix) were used for *in vivo* optogenetics. M1-ChR2 mice were stimulated at low intensity (~3 mW at the tip of the fiber), while S1-ChR2 and PV-ChR2 mice were stimulated at maximal intensity (~7.5 mW) for all behavioral tasks. Dual optogenetic mice (PV-NpHR and S1-ChR2) were stimulated at maximal intensity for both PV-NpHR (~3.5 mW) and S1-ChR2 (~7.5 mW). High intensity M1 stimulation was not used because it induced overt torso, limb, and facial movements, which interfered with licking, whereas low intensity S1 stimulation had no apparent effects on behavior. Stimulation intensity was kept consistent between mice in all three conditions by measuring the intensity prior to testing it with a PMD-100D optical power meter (Thorlabs).

Light was delivered to the cortical afferents and PV interneurons in the striatum through an optical fiber patchcord (Thorlabs; length 1 m; 0.50 nA; 200 µm diameter) connected to an optical cannula (described above) via a mating sleeve (Thorlabs). A small piece of heat shrink resistant tubing (Qualtek) was placed over the cannula during LED testing to prevent stray light during the task which may lead to texture illumination. A Pulse Pal [82] activated the LED current driver with 5 ms pulses at 25 Hz for two seconds beginning when the texture was presented. Corticostriatal neurons are capable of following this 25 Hz firing rate [83]

### Go/No-Go Tactile Discrimination Task

Head-fixed mice were trained to utilize their whiskers to discriminate between two textures presented to the whiskers on a motorized stage in a random order based on custom-written code in LabVIEW (National Instruments). This software controlled a linear stage and stepper motor similar to previously described [59,84]. The stepper motor rotated arms holding the two textures. Mice were trained to lick a piezo spout when presented with the “Go” texture (100 grit sandpaper; P100) and to withhold licking when presented with the “NoGo” texture (1200 grit sandpaper; P1200). A piezo film sensor that detected licks was connected to a solenoid-controlled water delivery spout. Additionally, a solenoid-controlled air spout was aimed at the contralateral mystacial pad. The texture task was carried out in a darkened room to minimize non-tactile cues. If mice correctly licked when presented with the Go texture, they were provided a small water reward (~5 µL). If mice incorrectly licked when presented with the NoGo texture, they were punished with a brief air puff. Sessions could be ended early if the mouse was no longer performing the task. Water was manually delivered (“autoreward”) by the experimenter following 15 consecutive trials without responding. If the mouse did not lick when water was present on the end of the spout following three autorewards, the session was ended.

Trials began with a 1 s pre-task interval, followed by texture movement toward the mice which was accompanied by a brief, cue tone (100 ms, 2930 Hz). Once the texture reached a set distance within reach of the whiskers, mice had a 2 s presentation time (PT) window to respond. A grace period was present during the initial 500 ms of the PT window where licks were not counted. Correctly licking in response to the Go texture (Hit) resulted in delivery of a water reward accompanied by a tone, while incorrectly licking in response to the NoGo texture (False Alarm; FA) resulted in a brief air puff (100-200 ms, 10-20 psi), a time-out period (7000-10000 ms), and an accompanying white noise. If no responding was detected to the Go or NoGo texture, the trial outcome was either a Miss or Correct Rejection (CR), respectively. During LED optogenetic stimulation, a 2 s train of 5 ms pulses at 25 Hz accompanied the PT window. Once mice either responded or 2 seconds had elapsed, the texture retreated to its original position where the next texture was rotated into position. Trials were separated by a 2 second intertrial interval.

Behavioral training lasted up to 3 weeks and mice were tested twice daily. Training proceeded in three general steps, but sessions could be added to ensure that mice adequately learned each step. (1) An initial shaping session acclimated mice to licking for a water reward. Water is automatically provided at the end of the PT window even if the mouse doesn’t lick. No texture is presented during this step. Each session consisted of 150 trials. (2) Only the Go texture is presented for two to three sessions, teaching mice to whisk against it and lick for a water reward. Each session consisted of 150 trials. (3) Both the Go and NoGo textures were presented at equal probabilities and no texture was presented for more than three consecutive trials. Mice were presented with interleaved Go and NoGo textures in a pseudorandom order. Mice were trained to whisk against the presented texture and respond appropriately. All sessions during the third step consisted of 127 trials. A subset of these sessions contained optogenetic stimulation. Sessions without optogenetic stimulation were interspersed to maintain behavioral performance. For single optogenetic stimulation testing (stimulation of S1 or M1 corticostriatal afferents or direct stimulation of PV interneurons), sessions were split into 50 baseline trials followed by 77 stimulation trials. For testing with dual optogenetic manipulation (stimulation of S1 corticostriatal afferents and inhibition of PV interneurons), sessions were divided into three blocks: 40 baseline trials, 40 S1-ChR2 stimulation trials and 47 S1-ChR2 and PV-NpHR stimulation trials. Stimulated and unstimulated data are from sessions where LED stimulation was administered. We used a block design to maximize the possibility of observing the effects of stimulation, and to reduce the number of behavioral sessions required by an interleaved stimulation design. Certain sessions were excluded from analysis due to poor performance under control conditions such as excessive impulsivity as indicated by an increase in FA rate or decreased motivation to complete the task which was indicated by an increase in Misses. Behavioral performance was further monitored by computing each mouse’s sensitivity (d’) score after each session. This score is derived from signal detection theory [85]. A d’ < 1 denoted non-expert performance, while a d’ > 1 denoted expert performance. If a mouse’s performance declined, it could be modulated by titrating water allowance or by training with more sessions without LED stimulation. At the end of behavioral testing, a single whisker trim control session was performed. The whiskers that were used to discriminate textures were unilaterally trimmed and performance was compared to the baseline period. Additionally, mice were tested while freely moving during open field (OF) and rotarod (RR) assays to examine if generalized motor activity increased due to optical stimulation as described in the Supplemental Information.

## Quantification and Statistical Analysis

### Analysis and Statistics

Behavioral performance was measured using four indices: Hit Rate, FA Rate, Sensitivity, and Bias. Hit Rate was calculated by the following equation: [Hit/(Hit+Miss)], where Hit is the number of correct Go trials and Miss is the number of incorrect Go trials. A similar formula was used to calculate FA Rate: [FA/(FA+CR)], where FA is the number of incorrect NoGo trials and CR is the number of correct NoGo trials. Sensitivity was calculated with the following equation: [(normalized inverse(Hit Rate)-normalized inverse(FA Rate))]. Bias was calculated with the following equation: [0.5*(normalized inverse(Hit Rate)+normalized inverse(FA Rate))]. Sensitivity illustrates how well a mouse can discriminate between two textures by comparing the normalized Hit and FA Rates. Bias is a measure that is independent of sensitivity and it illustrates a mouse’s overall responding, regardless of trial type.

Group data are presented as mean ± SEM. Statistics were calculated in MATLAB or SAS (SAS Institute). Data were analyzed and compared using paired or unpaired *t*-tests, one-way Analysis of Variance (ANOVA), or Repeated Measures ANOVA followed by post-hoc Bonferroni multiple comparison tests or paired contrasts respectively. In all cases, differences were considered significant at *p* < 0.05.

## Supplemental Material

**Figure S1:**
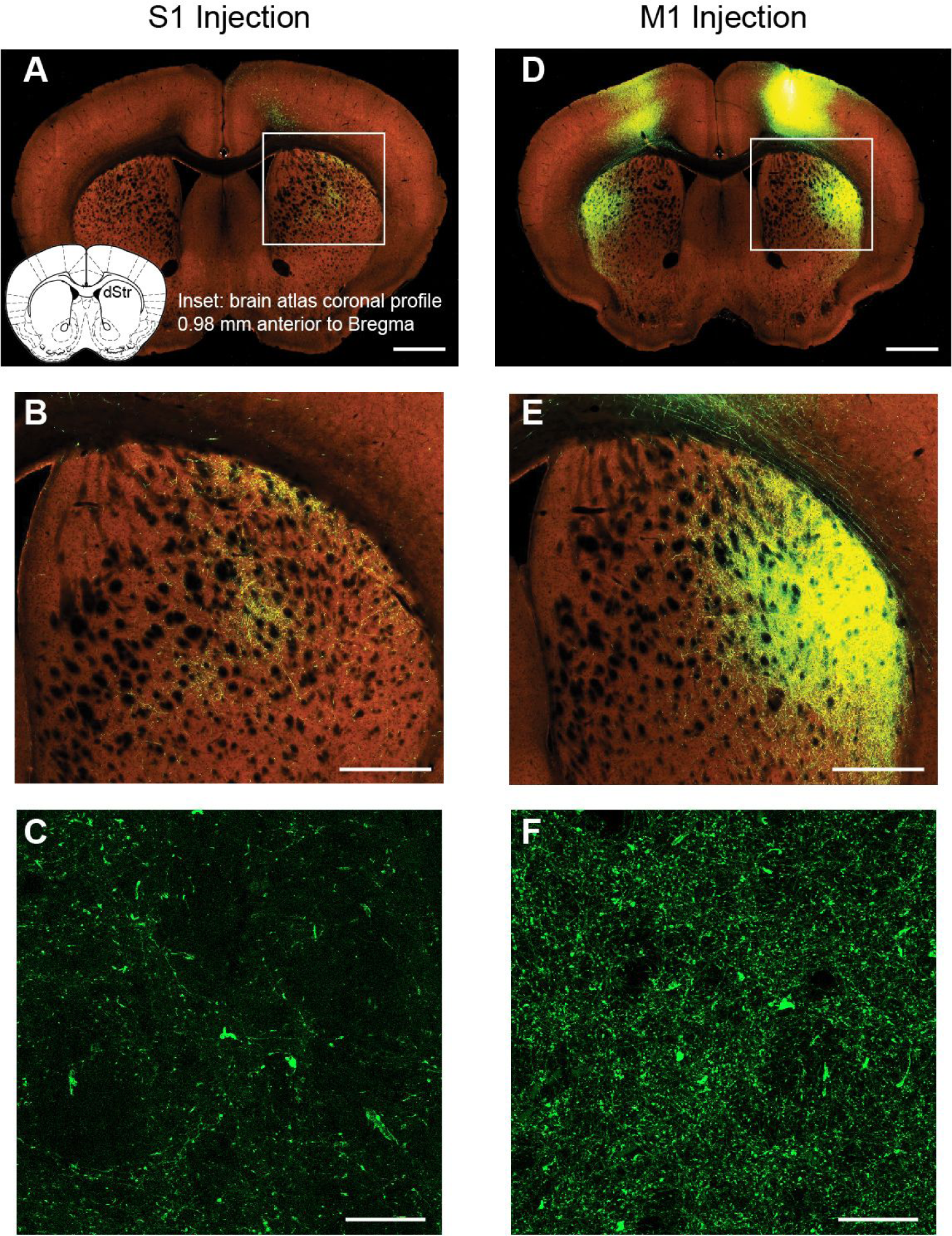
S1 and M1 have overlapping projection fields in dorsal anterior striatum. Related to Figures 1 and 2. **(A)** S1 barrel cortex injection of an enhanced green fluorescent protein adeno associated viral vector reveals corticostriatal fibers in dorsal anterior striatum. Inset shows a brain atlas coronal profile at 0.98 mm anterior to bregma which closely approximates the location of the fluorescent image [1]. **(B)** Higher magnification image of the inset indicated in **A** shows arborization of S1 corticostriatal axons throughout much of the central to lateral extent of dorsal anterior striatum. **(C)** Confocal maximal intensity projection of dorsal anterior striatum shows S1 corticostriatal fibers expressing ChR2-eYFP. **(D)** M1 corticostriatal fibers expressing EGFP in dorsal anterior striatum. **(E)** Higher magnification image of the inset in **D** shows arborization of M1 corticostriatal axons through a similar extent of striatum. **(F)** Confocal maximal intensity projection of dorsal anterior striatum reveals M1 corticostriatal afferents expressing ChR2-eYFP. **A**, **B**, **D** and **E** are from the Allen Mouse Brain Connectivity Atlas http://connectivity.brain-map.org/ [2]. Scale bars **A, D** 1 mm; **B, E** 500 µm; **C, F** 30 µm.

**Figure S2:**
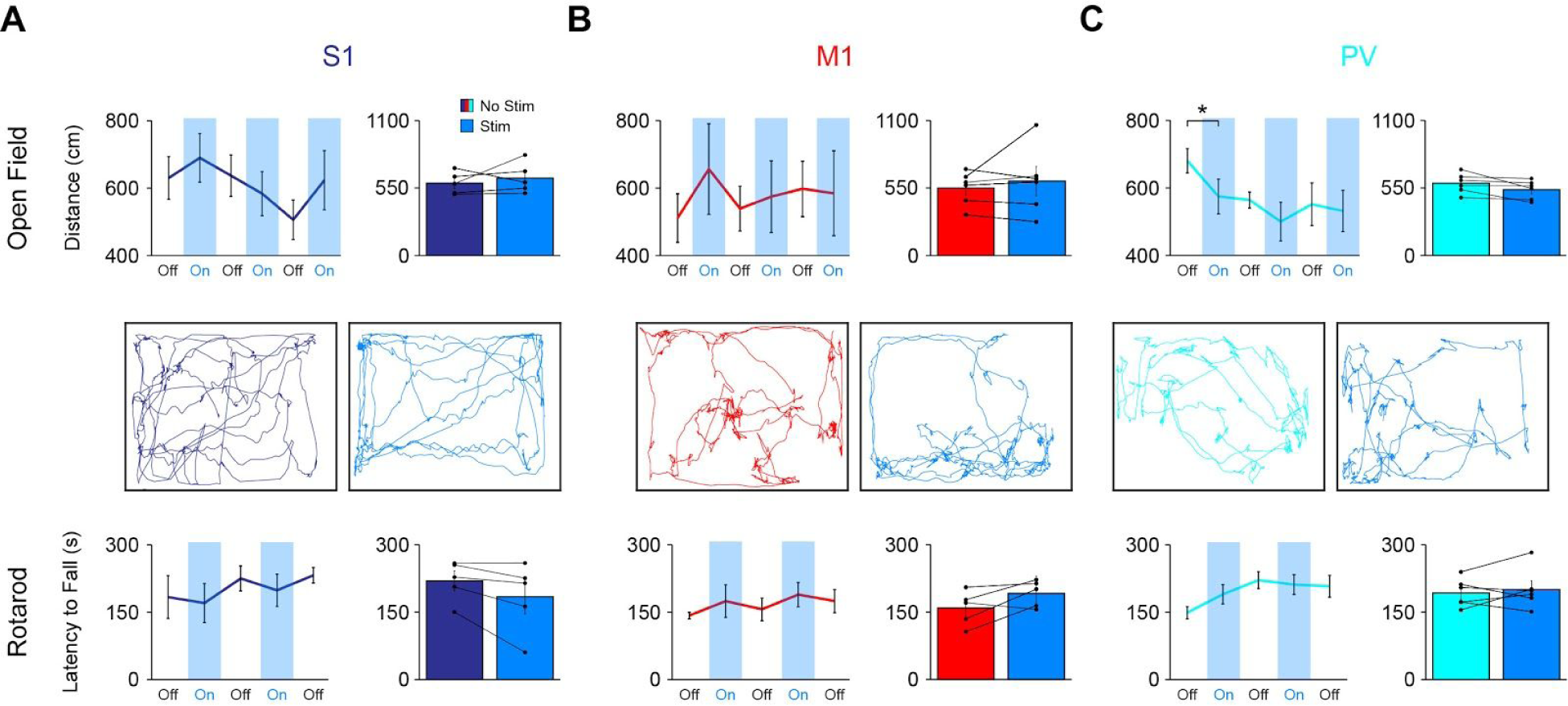
Stimulation of S1 or M1 corticostriatal afferents or PV striatal interneurons does not affect general motor behavior. Related to Figures 3 and 4. **(A)** *Top left*: Average total distance (± SEM) walked during 2 minute periods when S1 terminals (n = 5), **(B)** M1 terminals (n = 6), or **(C)** PV interneurons (n = 6) in the striatum were stimulated (blue) compared to not stimulated (white) in an open field task. *Top Right*: Grand mean of total distance traveled (± SEM) for **(A)** S1 terminals (n = 5), **(B)** M1 terminals (n = 6), or **(C)** PV interneurons (n = 6) in the striatum during baseline and stimulation periods. *Middle*: Representative tracking during 2 minute (*Left*) baseline and (*Right*) stimulation period in OF arena for **(A)** S1, **(B)** M1, and **(C)** PV conditions. *Bottom Left*: Average latency to fall (± SEM) when **(A)** S1 terminals (n = 5), **(B)** M1 terminals (n = 5), or **(C)** PV interneurons (n = 6) were either stimulated (blue) or not stimulated (white) on an accelerating rotarod task. Stimulation protocol was the same as the behavioral task and lasted up to 5 minutes. A 7 minute ITI followed each trial, allowing ample time for mice to recover, as well as precluding any lingering LED effects. Weight, ranging from 21.0g to 30.0g, showed no correlation with latency to fall using linear regression (r^2^ = 0.0113, n = 18). *Bottom Right*: Grand mean (± SEM) for **(A)** S1 terminals (n = 5), **(B)** M1 terminals (n = 5), or **(C)** PV interneurons during baseline and stimulation periods. No significant differences existed between the OF and RR baselines between the three groups (OF: (F_(2, 14)_ = 0.277, p = 0.762); RR: (F_(2, 13)_ = 3.186, p = 0.075)). Data shown as mean ± SEM. \**p* < 0.05.

**Figure S3:**
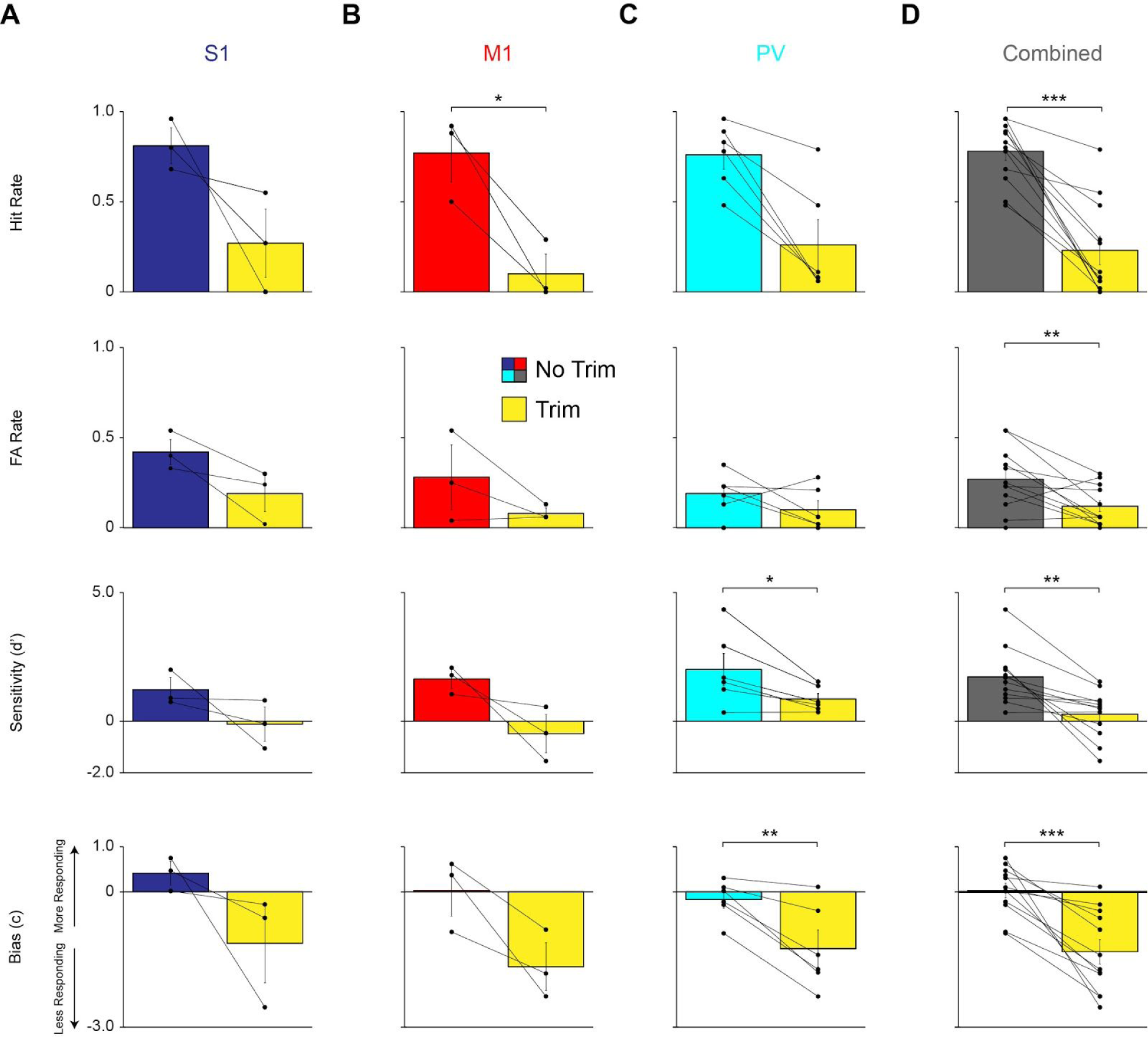
The texture discrimination task is dependent on tactile input. Related to Figures 3 and 4. Whiskers used during the sensorimotor choice task were unilaterally trimmed following 50 pre-trim trials. 100 post-trim trials were run and behavioral performance, based on our four parameters, was compared. Grand mean (± SEM) for pre- and post-whisker trim for **(A)** S1 (n = 3), **(B)** M1 (n = 3), **(C)** PV (n = 6), and **(D)** Combined (n = 12) conditions are shown based on each of the four parameters. The following are changes in the four parameters in the Combined condition: Hit Rate (0.78 ± 0.05 to 0.23 ± 0.08; p = 0.00002), FA Rate (0.27 ± 0.05 to 0.12 ± 0.03; p = 0.01), Sensitivity (1.72 ± 0.32 to 0.28 ± 0.28; p = 0.002), and Bias (0.03 ± 0.16 to −1.33 ± 0.27; p = 0.0003). \**p* < 0.05, \*\**p* < 0.01, \*\*\**p* < 0.001.

### Supplemental Experimental Procedures

#### Open Field Locomotor Behavior

Open field locomotor behavior was assessed by allowing mice to freely explore a 42 cm x 22 cm x 20 cm (LxWxH) polycarbonate laboratory arena (a rat cage with no bedding). It was placed inside a dark behavioral testing room. The cage was evenly illuminated using both red (Prophotonix) and infrared (Advanced Illumination) spotlights for automated mouse tracking. The optical fiber patchcord (Thorlabs) was connected to the mouse prior to being placed in the arena. LED stimulation with 5 ms pulses at 25 Hz occurred continuously during each 2 minute stimulation period. A video camera (Manta GigE, Allied Vision) mounted above the arena and Streampix 6 software (NorPix) were used to record the animal’s movements at a frame rate of 25 Hz with frame triggering controlled by a Pulse Pal (Open Ephys). The arena was cleaned with 70% ethanol between mice to eliminate sensory cues that might influence exploration and sanitized with 10% bleach each day.

Testing took two days to complete with a single session per day. On day one, mice were allowed to freely explore the arena for 30 minutes with the patchcord attached to the cannula. On day two, mice were habituated inside the arena for 10 minutes. Following the habituation period, the mice followed a block trial order consisting of 3 conditions (No Stim exploration, exploration during optogenetic stimulation, and freely moving intertrial interval) that lasted 2 minutes each. This block design was repeated three times. Videos were recorded and saved for each No Stim and Stim period. These videos were analyzed offline using a custom-written MATLAB (Mathworks) algorithm that segmented the mouse profile and calculated a centroid value (in pixels) for each frame and distance traveled was calculated. Counterbalancing was not needed as each Stim period was followed by a Freely Moving period that prevented carryover effects of stimulation. Both habituation periods minimized anxiety or exploratory effects.

#### Accelerating Rotarod Test

A rotarod (ENV-576M, Med Associates) was set to accelerate from 4 to 40 rpm over 5 minutes and. Trials ended when mice fell off the rod or after 5 minutes had elapsed. If a mouse stopped ambulating and instead clutched the rotarod, the trial was considered ended when the mouse made a complete revolution around the rod. Data were recorded as latency to fall or end of the trial. The entire apparatus was cleaned with 70% ethanol between mice and sanitized with 10% bleach following all sessions.

Testing took two days to complete with a single session per day. On day one, mice were habituated at a fixed speed of 4 rpm for 2 minutes followed by a 7 minute ITI. This was done 3 times for each mouse. On day two, stimulation was introduced using the parameters described above throughout the entire 5 minute trial unless the trial was ended by the mouse falling or clutching the rod. The first two trials were No Stim trials with the first trial not being used in the final analysis to allow the mouse to practice the task. After the first two No Stim trials, the next trial was a Stim trial that was followed by a No Stim trial. This order was repeated (No Stim, Stim, No Stim, Stim, No Stim) until two stimulation trials had been performed and the test ended on a No Stim trial. After each trial, mouse had a 7 minute ITI. Trials were counterbalanced when appropriate to ensure there were no carryover effects.

#### Histology

Following all testing, mice were deeply anesthetized with Ketamine-Xylazine (300 mg/kg Ketamine; 30 mg/kg Xylazine; i.p.) and transcardially perfused with PBS followed by 4% paraformaldehyde. The brain was carefully extracted and stored for 24 hours in 4% paraformaldehyde at 4°C. After, the brain was stored in 30% sucrose in PBS Azide at 4°C until the brain had sunk. The tissue was sectioned at 40 µm with a Thermo Fisher Scientific Shandon Cryotome FSE cryostat in the coronal plane. Sections were mounted on slides and coverslipped with Aqua-Mount (Thermo Scientific). Fluorescent photomicrographs were obtained using a Zeiss AxioObserver Z1 inverted fluorescence microscope for verification of injection site, cannula placement, and viral expression. Confocal photomicrographs were acquired using a Zeiss LSM 800 confocal laser scanning microscope. Confocal z stack images of corticostriatal fibers were acquired using a 40 x objective and 1 µm step size. Data were acquired and processed using Zeiss Zen software. Maximal intensity projections were created from stacks of 10 images and brightness and contrast adjusted equally on the entire image.

